# Compound- and sex-specific medial prefrontal cortex rewiring after prenatal THC and CBD exposure

**DOI:** 10.64898/2026.02.13.705680

**Authors:** Alba Cáceres-Rodríguez, Olivier Lassalle, Daniela Iezzi, Shuai Wang, Pascale Chavis, Olivier J. Manzoni

**Author notes:** Corresponding author: Olivier Manzoni, PhD., INMED, INSERM U901, Parc Scientifique de Luminy, BP13 – 13273 Marseille Cedex 09, France.

## Abstract

The growing perception of cannabinoids as benign has increased perinatal exposure to both Δ9-tetrahydrocannabinol (THC) and cannabidiol (CBD), yet their long-term effects on prefrontal circuitry remain incompletely defined. Here we used whole-cell and field electrophysiology in adult mice (P100-140) of both sexes to compare how in utero exposure to THC or CBD (GD5-18) remodels layer-5 pyramidal neurons in the medial prefrontal cortex (mPFC). We quantified intrinsic excitability, excitatory and inhibitory synaptic transmission, the net E/I ratio, spontaneous excitatory and inhibitory currents kinetics, and canonical forms of synaptic plasticity (endocannabinoid-mediated LTD and NMDA-dependent LTP). Both cannabinoids shifted the adult mPFC toward a more pro-excitatory state: the E/I ratio was altered in a compound- and sex-dependent manner, and endocannabinoid-mediated LTD was abolished across groups. THC recapitulated previously reported deficits, producing male-specific increases in intrinsic excitability and accelerating excitatory current kinetics in females, demonstrating a conserved cross-species vulnerability of prefrontal circuits to gestational THC. CBD produced distinct signatures: female offspring showed marked increases in spontaneous excitatory event frequency and amplitude, indicating large-scale reorganization of excitatory and inhibitory inputs, while male offspring exhibited a selective and profound impairment of LTP. The combination of universal eLTD loss and CBD-specific LTP impairment in males yields a bidirectional loss of plasticity: an emergent synaptic rigidity in which circuits are unable to potentiate or depress effectively. Together, these convergent and divergent effects establish that prenatal exposure to THC and CBD produces lasting, sex-dependent rewiring of mPFC circuitry. Our results caution that the non-intoxicating profile of CBD does not preclude durable developmental impact and underscore the importance of considering both compound identity and sex when assessing the neurodevelopmental risks of perinatal cannabinoid exposure.

## Introduction

The growing perception of cannabinoids as innocuous remedies has led to a surge in the use of both Δ^9^-tetrahydrocannabinol (THC) and the non-intoxicating cannabidiol (CBD) during the perinatal period. Due to their high lipophilicity, these compounds readily cross the placental barrier during gestation and sequester in breast milk during lactation ^1–5^. This pharmacokinetic profile explains how exogenous cannabinoids can interfere with critical endocannabinoid signaling throughout distinct stages of neurodevelopment, potentially leading to permanent neurobiological alterations ^6,7,8,9^.

Previous work established a robust blueprint for these disruptions. In rats, gestational THC exposure leads to sex-specific synaptic deficits in adult progeny ^3,6,8^, notably characterized by a profound impairment of endocannabinoid-mediated long-term depression (eLTD) in the medial prefrontal cortex (mPFC) -a primary hub for executive control and emotional regulation^10^. This deficit effectively stripped prefrontal synapses of their primary mechanism for activity-dependent depression^10^. Furthermore, the mPFC remains vulnerable postnatally, as isolated lactational exposure to cannabinoids via breast milk profoundly disrupts the maturation of prefrontal circuits and adolescent behavior^1,11,12,13^.

More recently, research has expanded to the non-intoxicating cannabinoid, cannabidiol (CBD). We and others established that gestational CBD exposure (E5–E18) in mice triggers sex-specific disruptions in early-life communication and the developmental trajectory of affective behaviors^13–15^. Mirroring these behavioral phenotypes, research in the insular cortex (IC), a structure critical for sensory processing, emotion regulation, and cognitive function, revealed that prenatal CBD induces a loss of functional distinction between IC territories^16,17^. This manifests as a dysregulated excitatory/inhibitory (E/I) balance and altered intrinsic membrane properties^16^.

Despite these advances, important questions remain regarding the conservation of these pathophysiological ’signatures’ across species and across different cannabinoid compounds. Whether gestational exposure to distinct cannabinoids produces conserved or compound-specific signatures of prefrontal dysfunction remains unresolved. To address this gap, we performed a systematic, sex-disaggregated comparison of in utero THC and CBD exposure using whole-cell and field electrophysiology in adult mice. We mapped intrinsic excitability, excitatory and inhibitory synaptic transmission, the net E/I ratio, synaptic currents’ kinetics, and canonical forms of synaptic plasticity (LTP and endocannabinoid-mediated LTD) in layer-5 mPFC pyramidal neurons and contrasted these profiles across sex and compound. Our results show that developmental cannabinoid exposure does not yield a single, uniform outcome: THC and CBD remodel mPFC circuits via distinct, sex-dependent trajectories that nevertheless converge on a shared loss of endocannabinoid-dependent LTD; CBD additionally produces a male-specific impairment of LTP, and both compounds drive compound- and sex-specific shifts in the E/I ratio. These convergent and divergent effects culminate in a persistent state of circuit dysregulation, indicating that cannabinoid identity and offspring sex critically determine the long-term impact of perinatal cannabinoid exposure on prefrontal cortical function.

## Materials and methods

### Animals

Male and female C57BL/6J mice (8–10 weeks old) were purchased from Charles River Laboratories and housed in standard wire-topped Plexiglas cages (42 cm 27 cm 14 cm) under temperature and humidity-controlled conditions (temperature 21 1°C, 60 10% relative humidity, and 12 h light/dark cycles). Food and water were available *ad libitum*. After one week of acclimation, female pairs were placed with a single male mouse in the late afternoon. The morning a vaginal plug was found was designated as gestational day 0 (GD0) and pregnant mice were housed individually. From GD5 to GD18, dams were injected subcutaneously (s.c.) daily with vehicle, 3 mg/kg of CBD^13–16^ or 3 mg/kg of THC^10,13^ (NIDA Drug Supply Program), dissolved in a vehicle consisting of Cremophor EL (Sigma-Aldrich), ethanol, and saline at 1:1:18 ratios, and administered at a volume of 4 mL/kg. Control dams (SHAM) were injected with the same volume of vehicle solution. This doses of CBD or THC reach the embryonic brain and induces behavioral changes in the offspring^10,14–16^. For each litter, the date of birth was designated as postnatal day (PND) 0. All procedures were performed in conformity with the European Communities Council Directive (86/609/EEC) and the United States NIH Guide for the Care and Use of Laboratory Animals. The French Ethical Committee authorized the project (APAFIS #49376).

### Slice Preparation

Adult male and female mice (PND 100–140) were deeply anesthetized with isoflurane and sacrificed according to institutional regulations. The brain was sliced (300 μm) in the coronal plane with a vibratome (Integraslice, Campden Instruments) in a sucrose-based solution at 4°C (in mM: 87 NaCl, 75 sucrose, 25 glucose, 2.5 KCl, 4 , 0.5, 2 , and 1.25 ) as described previously for the PFC^18–20^. Immediately after cutting, slices containing the PFC were stored for 30 minutes at 32°C in a low-calcium artificial CSF (aCSF) containing (in mM): 130 NaCl, 11 glucose, 2.5 KCl, 2.4 MgCl2, 1.2 CaCl2, 23 NaHCO3, and 1.2 NaH2PO4, equilibrated with 95% O_2_/CO_2_ /5%, and then kept at room temperature until the time of recording.

### Electrophysiology

During recording, slices were placed in the recording chamber and continuously perfused at 2 mL/min with warm (32°C) low- solution using the receptor blocker gabazine 10 µM (SR 95531 hydrobromide; Tocris). Pyramidal neurons were visualized under a differential interference contrast microscope using an upright microscope with infrared illumination (Olympus). Whole-cell patch-clamp recordings were made from the soma of layer V/VI pyramidal neurons in the prelimbic PFC^18,19^.

For current-clamp and voltage-clamp recordings, patch pipettes were filled with an intracellular solution containing (in mM): 145 K^+^ gluconate, 3 NaCl, 1 MgCl_2_, 1 EGTA, 0.3 CaCl_2_, 2 Na^2+^ ATP, 0.3 Na^+^ GTP, and 0.2 cAMP, buffered with 10 HEPES (pH 7.25, osmolarity 290–300 mOsm). Electrode resistance was 2–3 MΩ. Access resistance was monitored (< 25 MΩ) and the potential reference of the amplifier was adjusted to zero before breaking into the cell. Cells were held at –70 mV. Current-voltage (I-V) curves were made by a series of hyperpolarizing to depolarizing current steps immediately after breaking into the cell. Membrane capacitance (Cm) was estimated by integrating the capacitive current evoked by a −2 mV pulse, while membrane resistance was estimated from the I-V curve around resting membrane potential (RMP). RMP was measured immediately after whole-cell formation. The intrinsic excitability curve was made by measuring the number of action potentials elicited by depolarizing current steps of increasing amplitude, while rheobase was determined using a series of depolarizing 10 pA current steps. Data were recorded with an Axopatch-200B amplifier, low-pass filtered at 2 kHz, digitized (10 kHz, DigiData 1440A), and analyzed using Clampex 10.7 (Molecular Devices).

#### Postsynaptic Spontaneous Activity and Evoked Plasticity

Spontaneous activity was recorded to isolate AMPA, NMDA, and GABA currents^16,21^. <u>AMPA-sEPSCs</u>: Recorded at −70 mV using K-gluconate internal solution previously described with gabazine (10 µM). GABA-sEPSCs Recorded at -70mV using a high chloride internal solution (in mM: 140 KCl, 1.6 MgCl2, 2.5 MgATP, 0.5 NaGTP, 2 EGTA, 10 HEPES). The pH solution was adjusted to 7.25–7.3 and osmolarity to 280–300 mOsm. <u>GABA-sIPSCs</u> were recorded at − 70 mV in presence of 20 μM CNQX (6-Cyano-7-nitroquinoxaline-2,3-dione disodium, an AMPA receptor antagonist, Tocris) and L-APV 50 μM (DL-2-Amino-5-phosphonopentanoic acid, a selective NMDA receptor antagonist, Tocris). NMDA-sEPSCs: Recorded at +30 mV in the presence of 20 µM NBQX (2,3-dioxo-6-nitro-7-sulfamoyl-benzo[f]quinoxaline, Tocris) to block AMPA and kainite mediated currents and using a cesium-methanesulfonate internal solution (in mM: cesium-methanesulfonate (143), NaCl (5), MgCl2(1), EGTA (1), CaCl2 (0.3), Hepes (10), Na2ATP (2), NaGTP (0.3) and cAMP (0.2) (pH 7.3 and 290 mOsm).

#### Analysis

Spontaneous events were analyzed with Axograph X using a double exponential template: for AMPA-sEPSCs, rise = 0.5 ms and decay = 3 ms; for GABA-IPSCs, rise = 0.2 ms and decay = 10 ms; for NMDA-sEPSCs, rise = 3 ms and decay = 10 ms. Detection threshold was 3.5 times the baseline noise SD (amplitude threshold 7 pA). Total charge transfer was calculated by summing the charge transfer of every individual event in 6 min of acquisition^16,21^.

#### AMPA/NMDA Ratio

Measured on EPSCs evoked by electrical stimulation in layer 2/3 while recording layer 5 pyramidal neurons at +30 mV in Cs-methanesulfonate internal solution. The NMDA component was isolated by bath application of NBQX (20 μM) and the AMPA component obtained by digital subtraction^22,23^.

#### Extracellular Recordings

Field excitatory postsynaptic potentials (fEPSP) were recorded in layer 5 by electrical stimulation on layer 2/3 at 0.1 Hz. The glutamatergic nature was confirmed using CNQX (10 µM) at the end of each recording. Stimulus intensity was set at 60% of maximal intensity after performing an input-output curve.

#### LTP Induction

plasticity was induced by a Theta-burst stimulation (TBS) consisting of five trains of bursts (4 pulses at 100 Hz, 200-ms interval), repeated 4 times at 10 s intervals^24^.

#### LTD Induction

plasticity was induced by a 10 Hz stimulation for 10 min. Plasticity was calculated by comparing 10 min of baseline to the mean response 30 min post-induction^25^.

### Data Analysis and Statistics

Data were analyzed off-line with Clampfit 11.3 and AxoGraph 1.7.6. Graphs were generated with GraphPad Prism 10.4.1. Datasets were tested for normality (D’Agostino-Pearson and Shapiro-Wilk) and outliers (ROUT test) before running parametric or non-parametric tests. Statistical significance was assessed with two-way ANOVA followed by Sidak’s multiple comparison post-hoc tests. Quantitative data are presented as violin plots (median, 25th and 75^th^ quartile) with individual data points superimposed.

#### Synaptic Distribution and E/I Balance Analysis

Statistical analysis was performed in R (version 4.2.2) and GraphPad Prism 10.4.1. Cumulative distributions of excitatory and inhibitory charge transfer were compared using the Kolmogorov-Smirnov (KS) test. To evaluate the net functional balance, we calculated a net E/I Index (E/[E+I]). Because E and I current were recorded from independent populations of neurons, a nonparametric bootstrap was performed in R (version 4.2.2). For each group, the vectors of E and I value were resampled 10,000 times with replacement using the sample() function. Within each bootstrap iteration, we computed the mean E and mean I, using built-in R functions, and calculated the corresponding E/I index (E/(E+I)). This procedure yielded 10,000 bootstrap estimates of the E/I index per group. To quantify between-group differences, we computed the ΔE/I index by subtracting the bootstrap distributions (i.e., Group Sham minus Group CBD or THC). The proportion of ΔE/I values greater or less than zero was then used to assess significance. Following a standard bootstrap-based criterion, distributions in which more than 95% of ΔE/I values fell on one side of zero were interpreted as significant at p < 0.05.

To facilitate reading and interpretation, all statistical results are compiled in tables, while the figures display only the comparisons between male and female controls and the within-sex comparisons used to assess the effects of each compound.

## Results

In this study, we examined the long-term consequences of prenatal exposure to CBD and THC. To verify that gestation proceeded normally, several parameters were monitored throughout pregnancy. As shown in Table 1, pregnant dams exhibited comparable weight gain across groups, and pups were delivered around gestational day 19. Litter size and sex ratio were also unaffected. Together, these observations indicate that pregnancy progressed normally despite the daily subcutaneous cannabinoid treatment administered from gestational day 5 to 18.

### Prenatal cannabinoid exposure induces divergent biophysical signatures in mPFC pyramidal neurons

To assess how gestational cannabinoid exposure influences the fundamental biophysical properties of principal neurons in the medial prefrontal cortex (mPFC), we analyzed the passive membrane characteristics of Layer V pyramidal neurons in adult offspring (**Fig. 1**). These measurements indicate that THC reorganize neuronal membrane properties through distinct, sex-dependent mechanisms.

**Figure 1.**
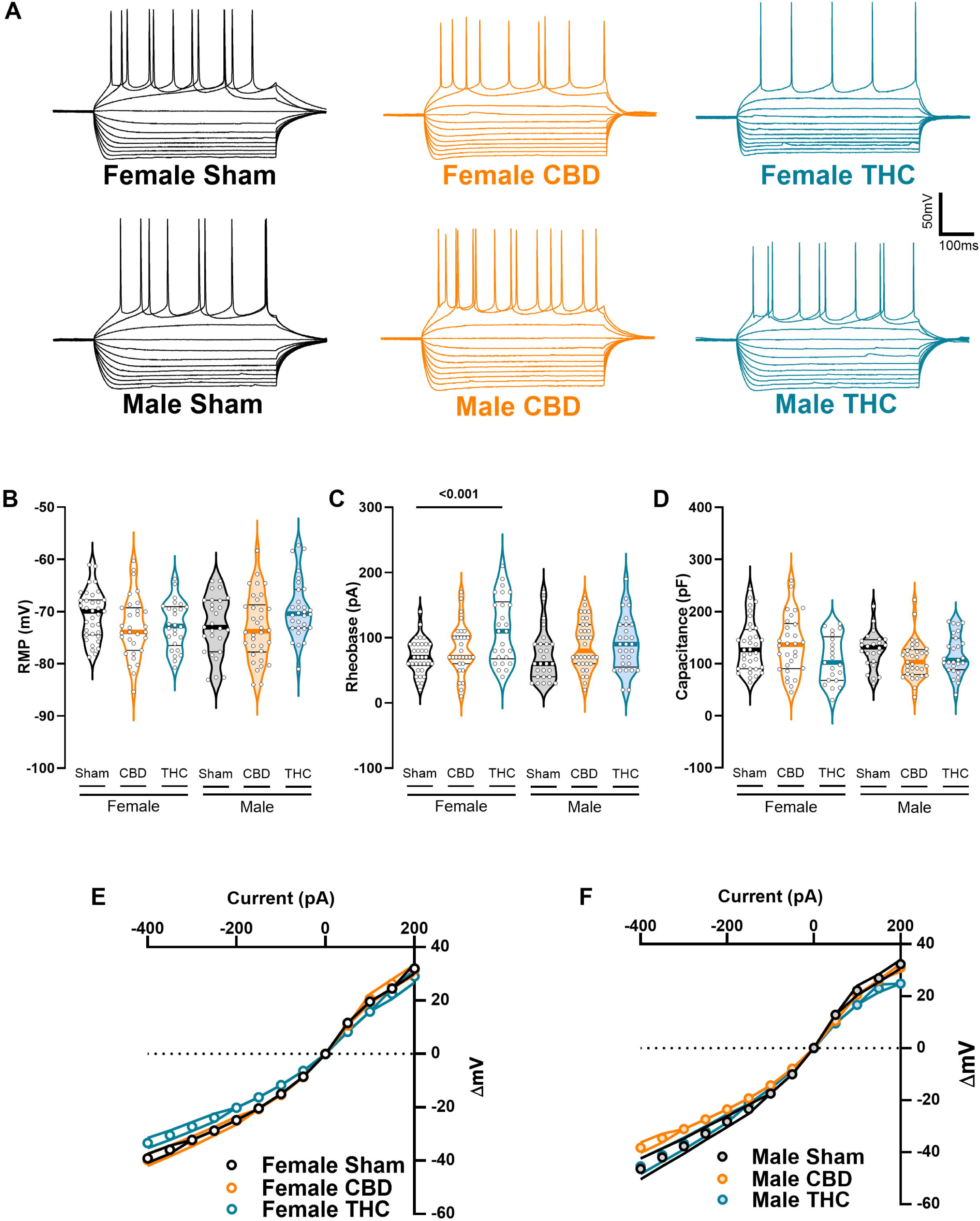
Prenatal cannabinoid exposure induces sex-specific changes in the passive properties of mPFC principal neurons: (A) Representative firing patterns evoked by hyperpolarizing and depolarizing current injections (500 ms; −400 to +150 pA in 50-pA steps) in Sham-, CBD-, and THC-exposed offspring of both sexes. Recordings were obtained from layer V pyramidal neurons of the medial prefrontal cortex (mPFC; Bregma +2.34). (B-D) Quantitative analysis of passive membrane properties of mPFC pyramidal neurons. (B) Resting membrane potential was not altered. (C) Progressive depolarizing current injections (10-pA steps) reveal an increased rheobase in females prenatally exposed to THC. (E) Capacitance, an indirect measure of cell size, remain similar across conditions. (E-F) No significant differences were observed in current–voltage (I–V) relationships of mPFC pyramidal neurons across sex or treatment. Responses were elicited by incremental current injections ranging from −400 to +200 pA in 50-pA steps. (B–D) Each dot represents a single neuron; data are presented as violin plots (median, and 25th–75th percentiles) and were analyzed using two-way ANOVA followed by Šídák’s multiple-comparison test. (E-F) Each dot represents the average response of a neuron at each current step; data are shown as mean ±SEM and analyzed using multiple Mann–Whitney U tests. Statistical significance (p-value < 0.05) are indicated in the graphs and in Table 1. Sample sizes (# neurons / # animals) were: females Sham (34/9), CBD (31/10), THC (26/9); males Sham (27/8), CBD (34/14), THC (30/8).

In both males and females, prenatal exposure to either CBD or THC did not alter the resting membrane potential (RMP), suggesting that the tonic membrane state of mPFC neurons remains preserved (**Fig. 1B**). In female offspring, however, THC (but not CBD) selectively increased the rheobase, indicating that a greater depolarizing current was required to reach firing threshold (**Fig. 1C**). Rheobase values in males were unaffected by either treatment (**Fig. 1C**, **Table 2**).

Across all treatment groups, membrane capacitance and the steady-state current–voltage (I–V) relationship remained stable (**Fig. 1D-F**, **Table 3**). These findings suggest that gestational cannabinoid exposure induces targeted modifications of specific biophysical features rather than broad alterations of the core resistive properties of the somatic membrane.

### Prenatal THC-induced hyperexcitability in males is driven by enhanced repetitive firing capacity independent of threshold shifts

We further examined the active discharge properties of mPFC neurons by assessing the number of evoked action potentials in response to increasing depolarizing current steps (**Fig. 2**, **Table 2 and 4**). These analyses revealed that the effect of prenatal THC on mPFC pyramidal neurons’ excitability is confined to male offspring, consistent with our previous observations in rats^10^. Female mPFC neurons showed stable intrinsic excitability and firing rates across all treatment groups (**Fig. 2A**). In contrast, THC-exposed males exhibited a pronounced increase in intrinsic excitability, generating more action potentials in response to depolarizing current injections, particularly at higher stimulus intensities (from 350pA; **Fig. 2B, D**). Importantly, this hyperexcitability did not arise from alterations in the fundamental mechanism initiating action potentials, as AP threshold values were unchanged across sex and treatment conditions (**Fig. 2C**).

**Figure 2.**
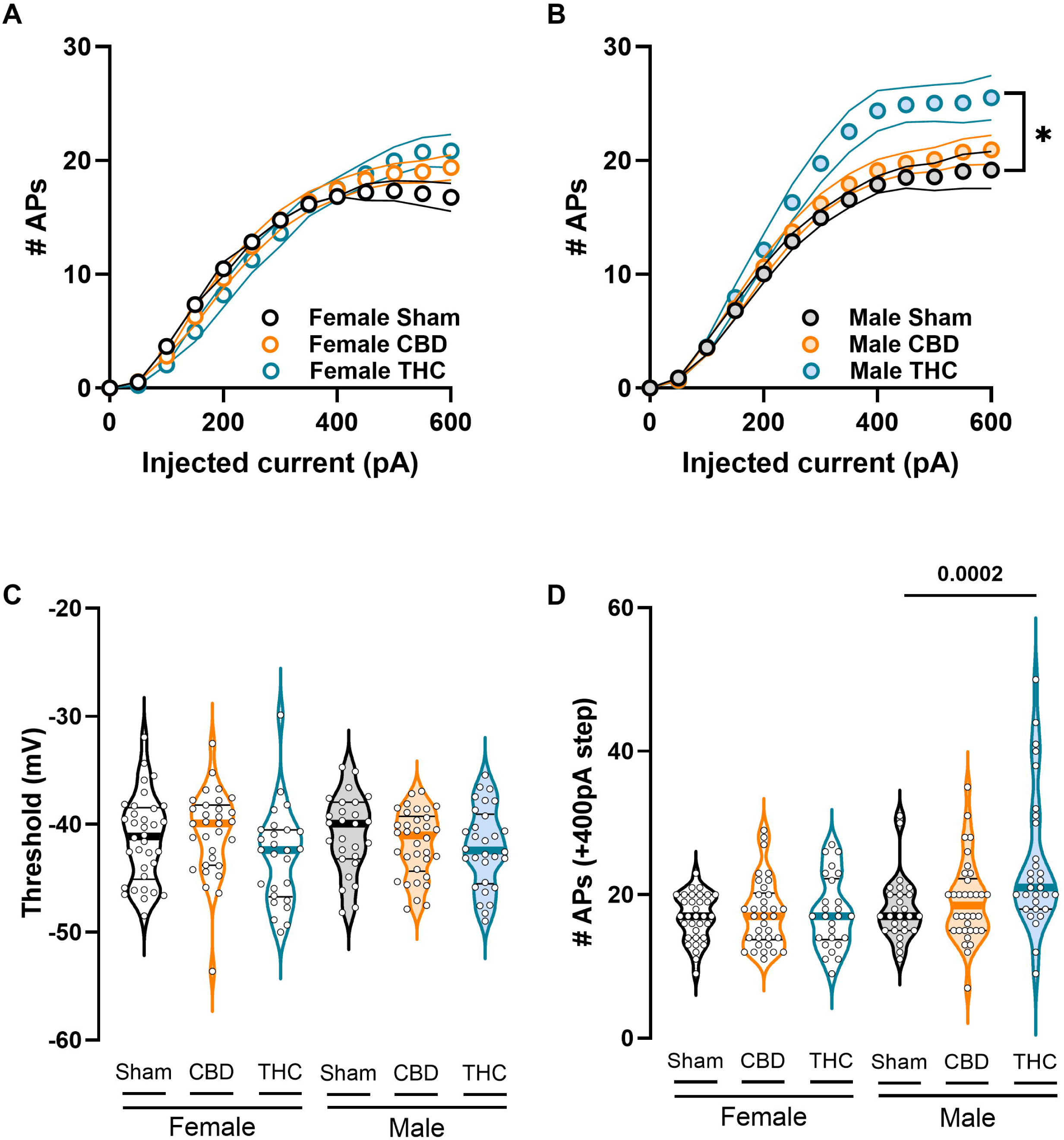
THC-exposed male progeny displays increased intrinsic neuronal excitability in adulthood. (A–B) Number of evoked action potentials in response to increasing depolarizing current injections. (A) Female offspring display comparable intrinsic excitability across experimental conditions. (B) Progressive depolarizing current injections (50-pA steps) reveal increased firing in THC-exposed males at supraphysiological stimulation intensities. (C) Despite this increase in firing, the action potential threshold does not differ across sex or treatment. (D) Representative example showing the number of spikes elicited by a +400-pA depolarizing current step, illustrating enhanced firing in THC-exposed males. (A–B) Each dot represents the group mean value at the corresponding current step; data are presented as mean ± SEM in XY plots and were analyzed using multiple measures Mann-Whitney Test. (C–D) Each dot represents a single neuron; data are shown as violin plots (median, and 25th–75th percentiles) and were analyzed using two-way ANOVA followed by Šídák’s multiple-comparison test. Only statistically significant differences (p < 0.05) are indicated in the graphs. Sample sizes (# neurons / # animals) were: females Sham (34/9), CBD (31/10), THC (26/9); males Sham (27/8), CBD (34/14), THC (30/8).

### Prenatal cannabinoids induce divergent remodeling of spontaneous excitatory synaptic transmission

To determine whether the observed changes in intrinsic excitability were accompanied by alterations in synaptic input, we recorded spontaneous excitatory postsynaptic currents (sEPSCs) from layer V pyramidal neurons (**Fig. 3A–B**). These measurements revealed a sex-dependent reorganization of excitatory drive following gestational cannabinoid exposure.

**Figure 3.**
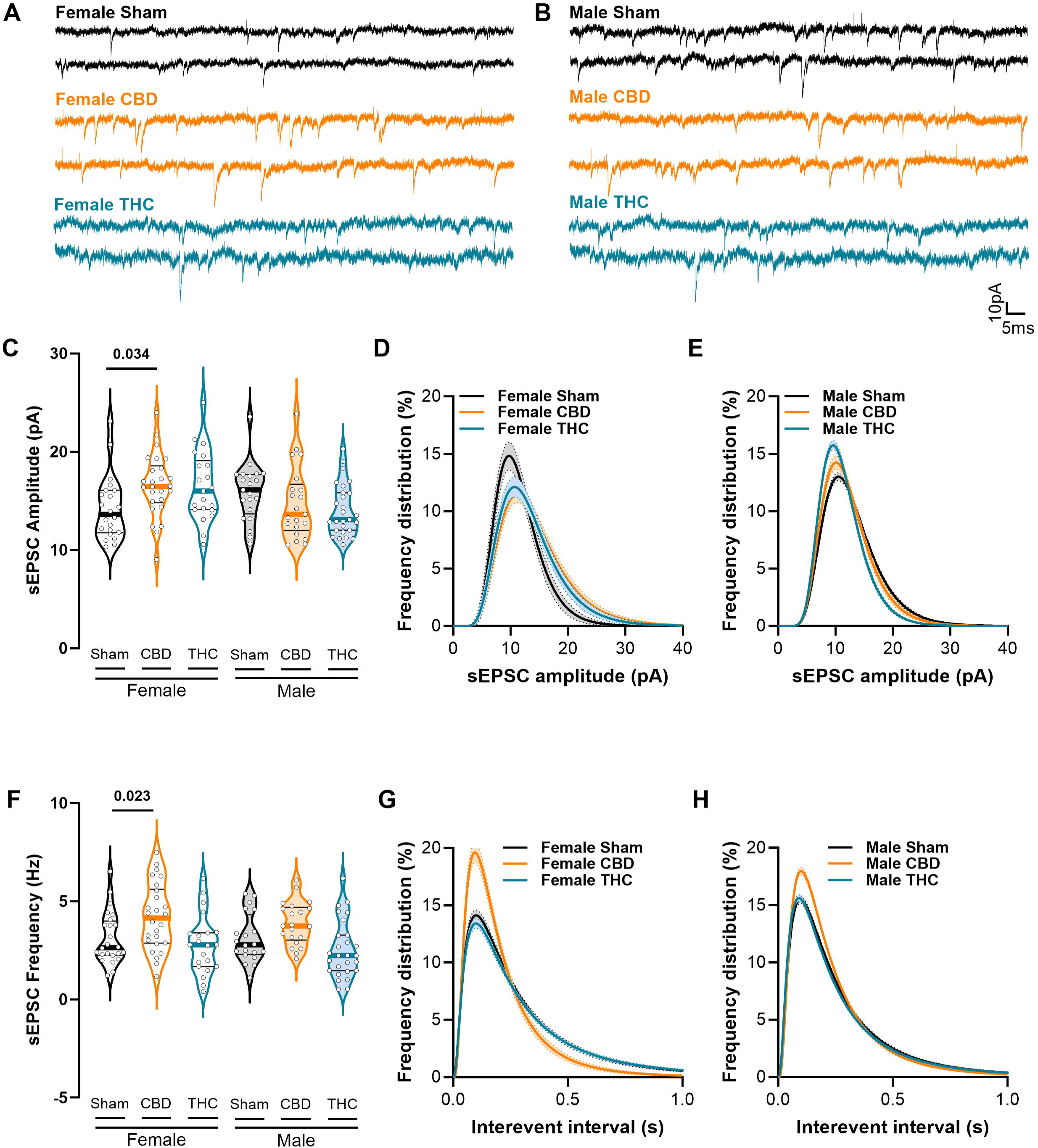
Prenatal cannabinoid exposure alters excitatory synaptic transmission in female progeny: (A–B) Representative traces of spontaneous excitatory postsynaptic currents (sEPSCs) recorded at −70 mV in presence of gabazine (10µM) from Sham-, CBD-, and THC-exposed female and male offspring (scale bar 10pA/5ms). (C) On average, CBD-exposed females exhibit larger sEPSC amplitudes. (D) Log-normal distribution fitting (± CI) reveals a higher proportion of large-amplitude events in cannabinoid-treated females, whereas (E) cannabinoid-treated males display a shift toward smaller-amplitude events. (F) sEPSC frequency is increased in CBD-exposed females compared with Sham and THC groups. (G) Consistent with this, inter-event intervals are reduced in CBD-exposed females relative to Sham and THC females. (H) A higher proportion of CBD-exposed males exhibit shorter inter-event intervals, indicating increased event frequency. (C, F) Data are presented as violin plots (median, and 25th–75th percentiles) and were analyzed using two-way ANOVA followed by Šídák’s correction for multiple comparisons. (D, E, G, H) Data were analyzed using log-normal curve fitting with confidence intervals (± CI). Only statistically significant differences (p < 0.05) are indicated in the graphs additional statistical information can be found in Table 5. Sample sizes (# neurons / # animals) were females Sham (24/9), CBD (26/12), THC (21/9); males Sham (21/8), CBD (21/13), THC (27/8).

In female offspring, prenatal CBD exposure markedly enhanced excitatory transmission, reflected by increases in both mean sEPSC amplitude (**Fig. 3C-D**) and event frequency (**Fig. 3F-G**). Log-normal distribution analysis showed a pronounced rightward shift, indicating a greater proportion of large-amplitude events in CBD-treated females (**Fig. 3D**). Consistent with this, inter-event intervals were significantly shortened (**Fig. 3G**), confirming an elevated rate of glutamate release. In contrast, although THC-exposed females exhibited a shift toward larger-amplitude events (**Fig. 3D**), this change did not translate into an increase in average sEPSC amplitude (**Fig. 3C**), and their event frequency remained comparable to Sham controls (**Fig. 3 F-G**).

Male offspring displayed a distinct synaptic phenotype. Both THC and CBD exposure increased the proportion of small-amplitude sEPSCs (**Fig. 3E**), but these shifts were insufficient to alter mean event amplitude. CBD-exposed males also showed a higher proportion of neurons with short inter-event intervals (**Fig. 3H**), yet this did not significantly affect their average event frequency (**Fig. 3F**).

### Sex-specific reorganization of the inhibitory synaptic landscape

We subsequently characterized spontaneous inhibitory postsynaptic currents (sIPSCs) in our various treatment groups, to determine if the GABAergic inhibitory drive within the mPFC microcircuitry was subject to a concomitant reorganization (**Fig. 4 and Table 6**).

**Figure 4.**
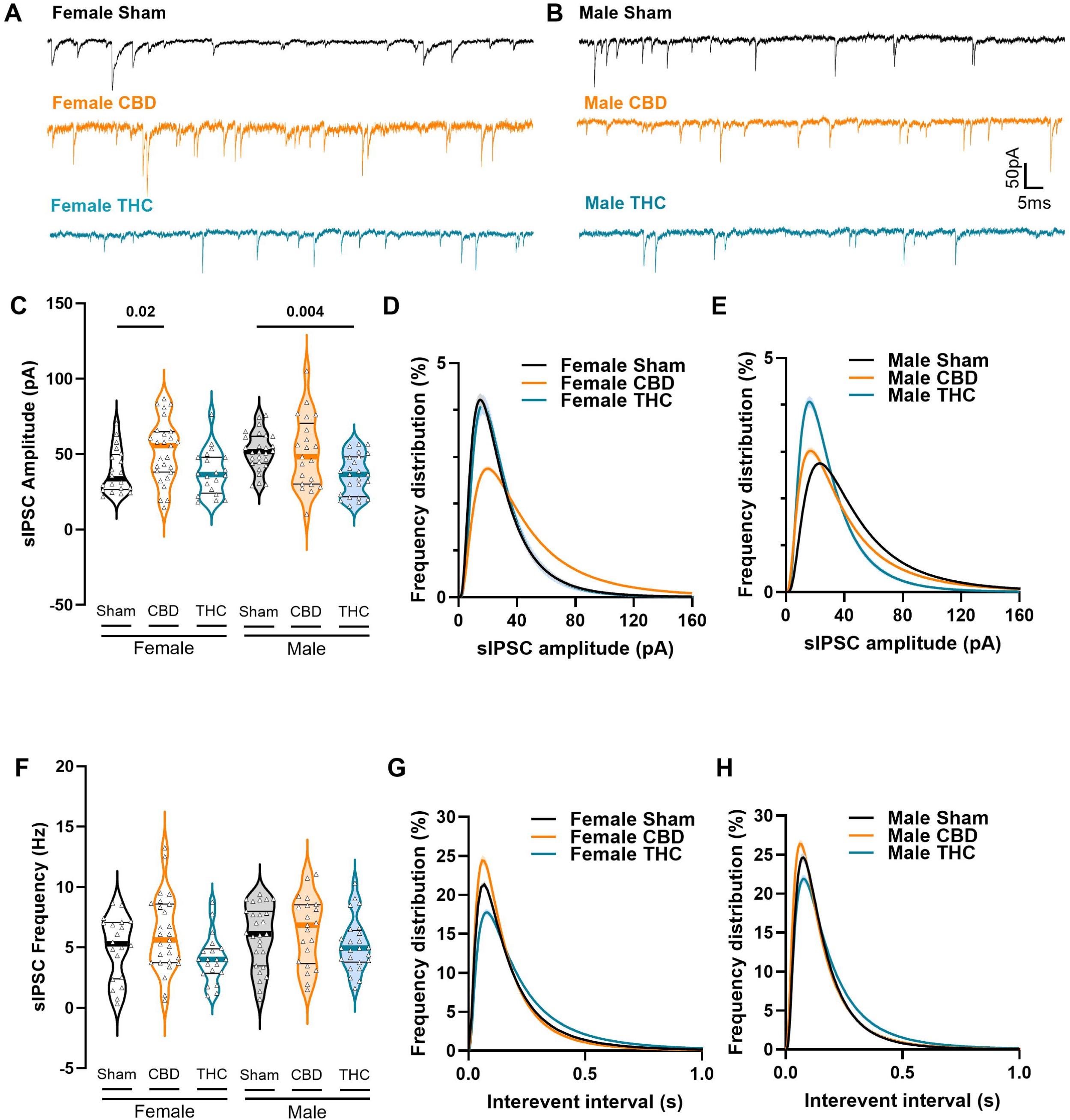
Prenatal cannabinoid exposure differentially alters inhibitory synaptic transmission in female and male progeny: (A–B) Representative traces of spontaneous inhibitory postsynaptic currents (sIPSCs) recorded at −70 mV from Sham-, CBD-, and THC-exposed female and male offspring in the presence of APV and CNQX (10 µM), (scale bars: 50 pA, 5 ms). (C) On average, CBD-exposed females exhibit larger sIPSC amplitudes. (D) Log-normal distribution fitting (± CI) reveals a higher proportion of large-amplitude events in CBD-treated females, whereas (E) cannabinoid-treated males display a shift toward smaller-amplitude events. (F) Mean sIPSC frequency does not change in cannabinoid-treated females. (G) However in the distributions, inter-event intervals are reduced in CBD-exposed females relative to Sham and THC females. (H) Conversely, THC-exposed males display a higher proportion of longer inter-event intervals, indicative of reduced event frequency. (C, F) Data are presented as violin plots (median, and 25th–75th percentiles) and were analyzed using two-way ANOVA followed by Šídák’s correction for multiple comparisons. (D, E, G, H) The graphs display the relative distribution of all events in 6min recording, these data were analyzed using log-normal curve fitting with confidence intervals (± CI). Only statistically significant differences (p < 0.05) are indicated in the graphs. Further statistical details can be found on Table 6. Sample sizes (# neurons / # animals) were females Sham (20/6), CBD (27/7), THC (21/5); males Sham (27/7), CBD (21/7), THC (24/5).

Consistent with the excitatory findings, CBD-exposed females exhibited a robust gain in inhibitory transmission, showing increased sIPSC mean amplitudes (**Fig. 4C**) along a significant enrichment of large-amplitude inhibitory events in this group (**Fig. 4D**). sIPSC frequency remained unchanged (**Fig. 4F-G**).

Conversely, male progeny showed a general weakening of inhibitory synaptic strength. THC males display decreased mean amplitude (**Fig. 4C**). Both THC and CBD exposure resulted in a shift toward smaller sIPSC amplitudes (**Fig. 4E**). sIPSC frequency was unchanged in both groups, as observed in females (**Fig. 4F-H**).

THC-exposed males further displayed an increase in the proportion of longer inter-event intervals associated with a decrease, suggesting smaller frequency of GABA-A-R activation (**Fig. 4H**).

Collectively, these data (**Fig. 3 & 4**) reveal a sexually divergent synaptic signature: female CBD exposure induces an overall potentiation of synaptic inputs, while female THC distribution data reports an overall decrease in inhibitory transmission due to the extension of the inter-event interval of GABAergic currents. In parallel, male THC together with male CBD exposure results in an attenuation of the total excitatory and inhibitory synaptic drive.

### Gestational cannabinoids alter the temporal kinetics of synaptic inputs

Beyond frequency and amplitude, the timing of receptor activation and deactivation plays a critical role in shaping synaptic integration. To determine whether gestational cannabinoids also influence this temporal dimension of synaptic signaling, we examined the kinetics of spontaneous synaptic currents and found significant alterations in receptor-mediated timing (**Fig. 5**, **Table 5-6**). Across sexes, prenatal THC exposure produced markedly faster sEPSC rise times (**Fig. 5B**), while decay times remained unchanged (**Fig. 5C**). This acceleration of the rising phase is most consistent with a shift in AMPA receptor subunit composition—potentially favoring subunits with faster activation kinetics—although changes in synaptic microarchitecture or a redistribution of excitatory inputs toward more proximal dendritic compartments may also contribute.

**Figure 5.**
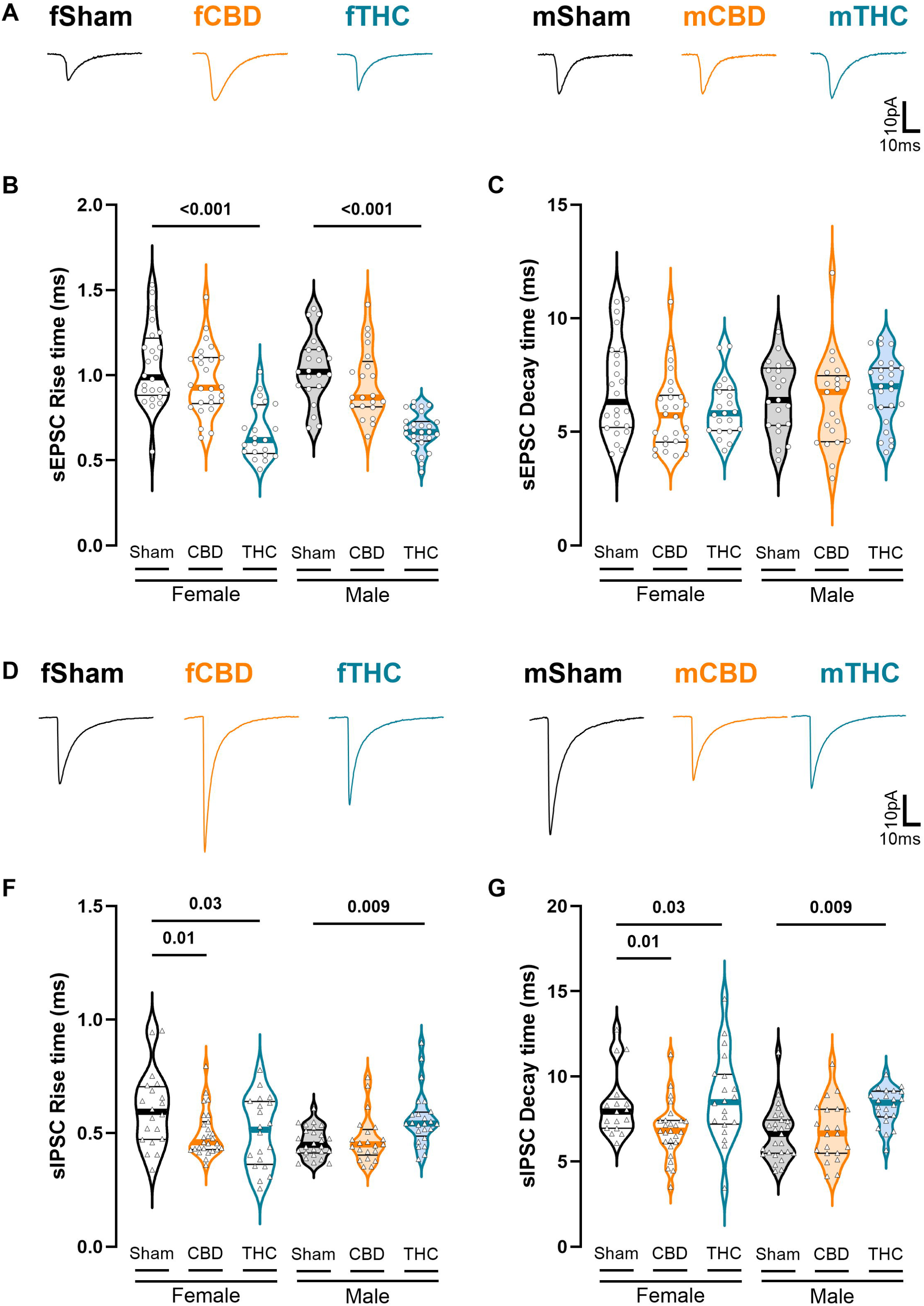
Receptor kinetics of excitatory and inhibitory synaptic transmission are differentially modified in a sex- and treatment-dependent manner: (A) Representative traces of sEPSC currents in each treatment group (B) Regardless of sex, THC-exposed progeny exhibits faster rise times of excitatory postsynaptic currents. (C) Decay times of excitatory postsynaptic currents remain unchanged across sex and treatment. (D) Representation on mean sIPSC currents per sex and treatment. (E) Compared with controls, inhibitory postsynaptic currents display faster rise times in cannabinoid-treated females, whereas cannabinoid-treated males exhibit slower rise times. (E) For inhibitory current decay kinetics, CBD-exposed females show faster decay times, while THC-exposed females exhibit prolonged decays; THC-exposed males also display increased decay times. (B, C, E, F) Data are presented as box-and-whisker plots as violin plots (median, and 25th–75th percentiles) and were analyzed using two-way ANOVA followed by Šídák’s correction for multiple comparisons. Only statistically significant differences (p < 0.05) are indicated in the graphs.

Inhibitory kinetics exhibited a more heterogeneous pattern. In females, both THC and CBD exposure accelerated sIPSC rise times (**Fig. 5E**). However, their effects on decay kinetics diverged: CBD-exposed females showed faster decay, whereas THC significantly prolonged inhibitory decay (**Fig. 5F**). In males, CBD had no discernable effects while THC generally slowed inhibitory signaling, with THC-treated males displaying increased rise and decay times (**Fig. 5E-F**). Together, these findings indicate that the temporal precision of inhibitory transmission is differentially “tuned” by the specific cannabinoid and the sex of the offspring, adding an additional layer of complexity to the synaptic reorganization induced by gestational exposure.

### Prenatal Cannabinoid Exposure Reorganizes mPFC E/I Balance

We assessed the functional impact of prenatal CBD and THC exposure on mPFC synaptic processing by quantifying the net excitatory–inhibitory (E/I) balance in male and female offspring (**Fig. 6**, **Table 7**). To integrate the previously observed changes in synaptic amplitude and frequency, we first analyzed the cumulative distributions of total excitatory and inhibitory charge transfer for each group.

**Figure 6.**
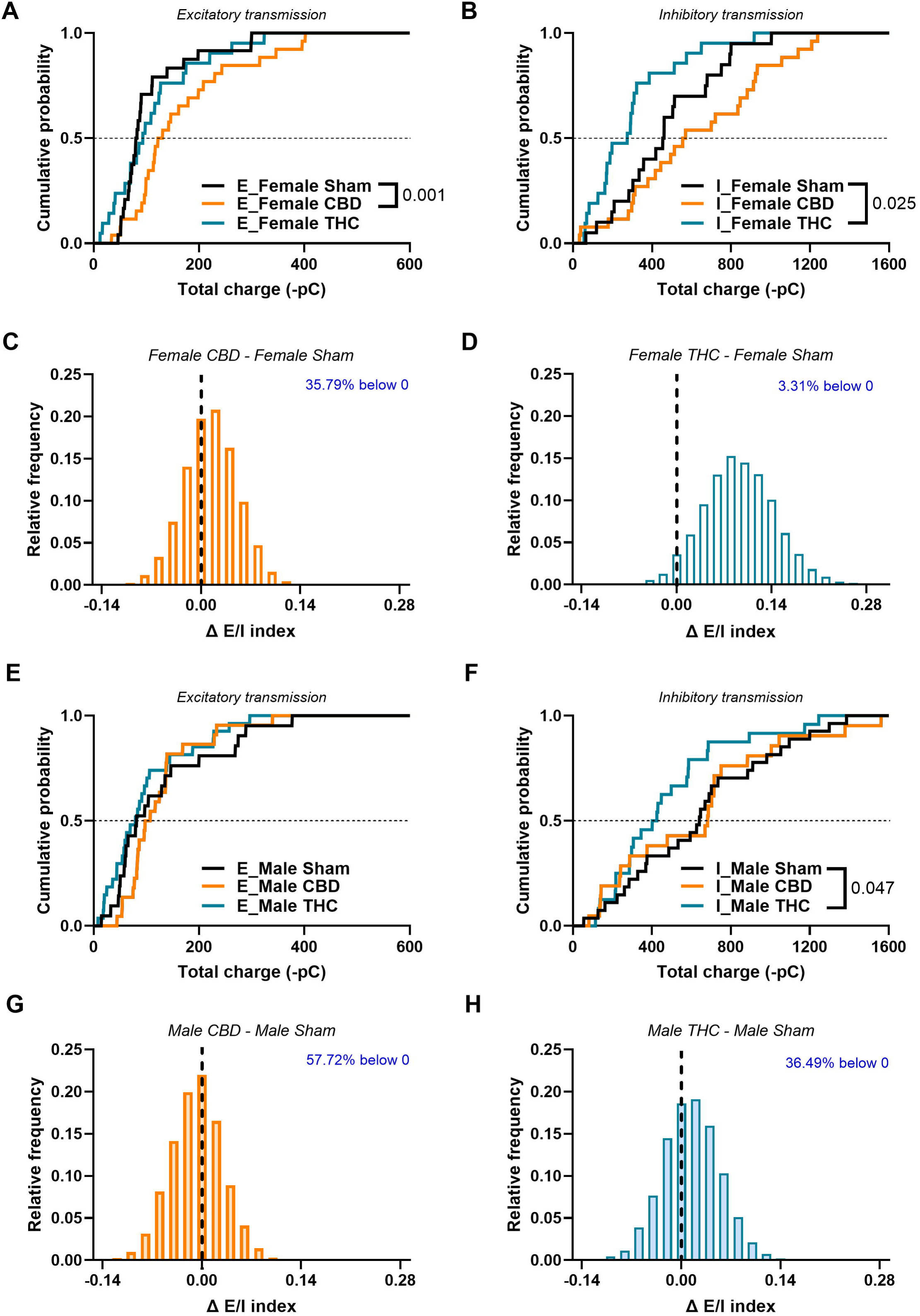
Prenatal cannabinoid exposure alters the excitatory/inhibitory balance. (A, E) Total charge transferred by AMPA-mediated sEPSCs and (B, F) GABA-mediated sIPSCs measured over a 6-min recording period across sexes and treatments. (A) CBD-exposed females show increased quantal excitatory transmission. (B) THC-exposed females display reduced inhibitory charge transfer compared with Sham controls. (C, D, G, H) Distribution plots of the difference between the bootstrapped E/I index of each cannabinoid-treated group and its corresponding Sham group. Distributions in which more than 95% of ΔE/I values fall on one side of zero are interpreted as significantly different (p < 0.05), indicating an E/I imbalance. (C) CBD-treated females show no change in E/I balance. (D) THC-treated females exhibit an increased E/I index driven by reduced inhibitory tone. (E) Total excitatory transmission remains unchanged in males, whereas (F) THC-exposed males show a marked reduction in inhibitory charge transfer. These modest synaptic changes account for the stability of the E/I index in cannabinoid-treated male offspring. (G, H) Neither CBD- nor THC-exposed males differ from Sham in their E/I index distributions. (A, B, E, F) Cumulative frequency distributions of total charge transfer for sEPSCs (excitatory) and sIPSCs (inhibitory) were obtained from the neurons recorded in Figures 4 and 5. (C, D, G, H) Relative distribution plots show the difference between the bootstrapped E/I index of each treated group and its corresponding Sham group. Only statistically significant differences (p < 0.05) are indicated in the graphs.

In females, the data reveal a cannabinoid-specific reorganization of synaptic inputs. Prenatal CBD exposure produced a marked rightward shift in the cumulative distribution of excitatory charge transfer (**Fig. 6A**, D = 0.555, p = 0.001), accompanied by a qualitative rightward trend in inhibitory charge transfer relative to Sham controls (D = 0.357, p = 0.106; **Fig. 6B**). This pattern suggests that the increase in excitatory drive may be partially compensated by a parallel enhancement of inhibitory input. In contrast, prenatal THC exposure elicited only a qualitative rightward shift in excitation (D = 0.238, p = 0.549; **Fig. 6A**), but this was paired with a very large leftward shift in inhibitory charge transfer (D = 0.462, p = 0.025; **Fig. 6B**, Table 7), suggesting a net reduction in inhibitory drive.

Male offspring displayed a different pattern of vulnerability. CBD exposure did not significantly alter excitatory (**Fig. 6E**, D = 0.297, p = 0.301) or inhibitory charge distributions (**Fig. 6F**, D =0.164, p =0.908; **Fig. 6E-F**), indicating relative synaptic stability. In contrast, THC-exposed males exhibited a small leftward shift in inhibitory charge transfer (D = 0.384, p = 0.047) while excitatory charge transfer remained unchanged (D = 0.201, p = 0.726; **Fig. 6 E-F**).

Group-level comparisons of the E/(E + I) index were obtained using a bootstrap-based estimation procedure (see Methods). Statistical significance was defined by the consistency of ΔE/I values relative to zero, with contrasts considered significant when more than 95% of bootstrap samples fell on the same side of zero (p < 0.05). As shown in **Fig. 6D**, this analysis revealed that THC-exposed females exhibit a significantly higher E/I index than Female Sham controls. In contrast, CBD-exposed females showed no net change in E/I balance, as the increase in excitatory drive was offset by a parallel tendency toward enhanced inhibitory input (**Fig. 6C**).

In males, neither CBD nor THC exposure produced significant changes in the E/I index (**Fig. 6G-H**), consistent with the relative stability observed in their underlying charge-transfer distributions.

Together, these findings demonstrate that gestational cannabinoid exposure reshapes the synaptic E/I landscape in a sex- and compound-specific manner: females show marked elevations in the E/I index—most prominently after THC exposure—whereas males display no significant changes in E/I ratio despite THC-induced alterations in inhibitory charge transfer.

### Prenatal CBD and THC Exposure Abolish eLTD in Mouse Progeny

We next examined how prenatal cannabinoid exposure affects endocannabinoid-mediated long-term depression (eLTD), a key form of synaptic flexibility in the mPFC and a mechanism we previously identified as vulnerable to prenatal THC exposure in rats^10,12^. In Sham-exposed offspring of both sexes, low-frequency stimulation (10 Hz, 10 min) reliably induced robust eLTD, reflected by a sustained reduction in fEPSP amplitude relative to baseline (**Fig. 7**). Strikingly, this form of plasticity was entirely absent in adult mice prenatally exposed to either CBD or THC. In both females (**Fig. 7A–B**) and males (**Fig. 7C–D**), fEPSP amplitudes failed to depress following induction and instead remained at or above baseline throughout the recording period.

**Figure 7.**
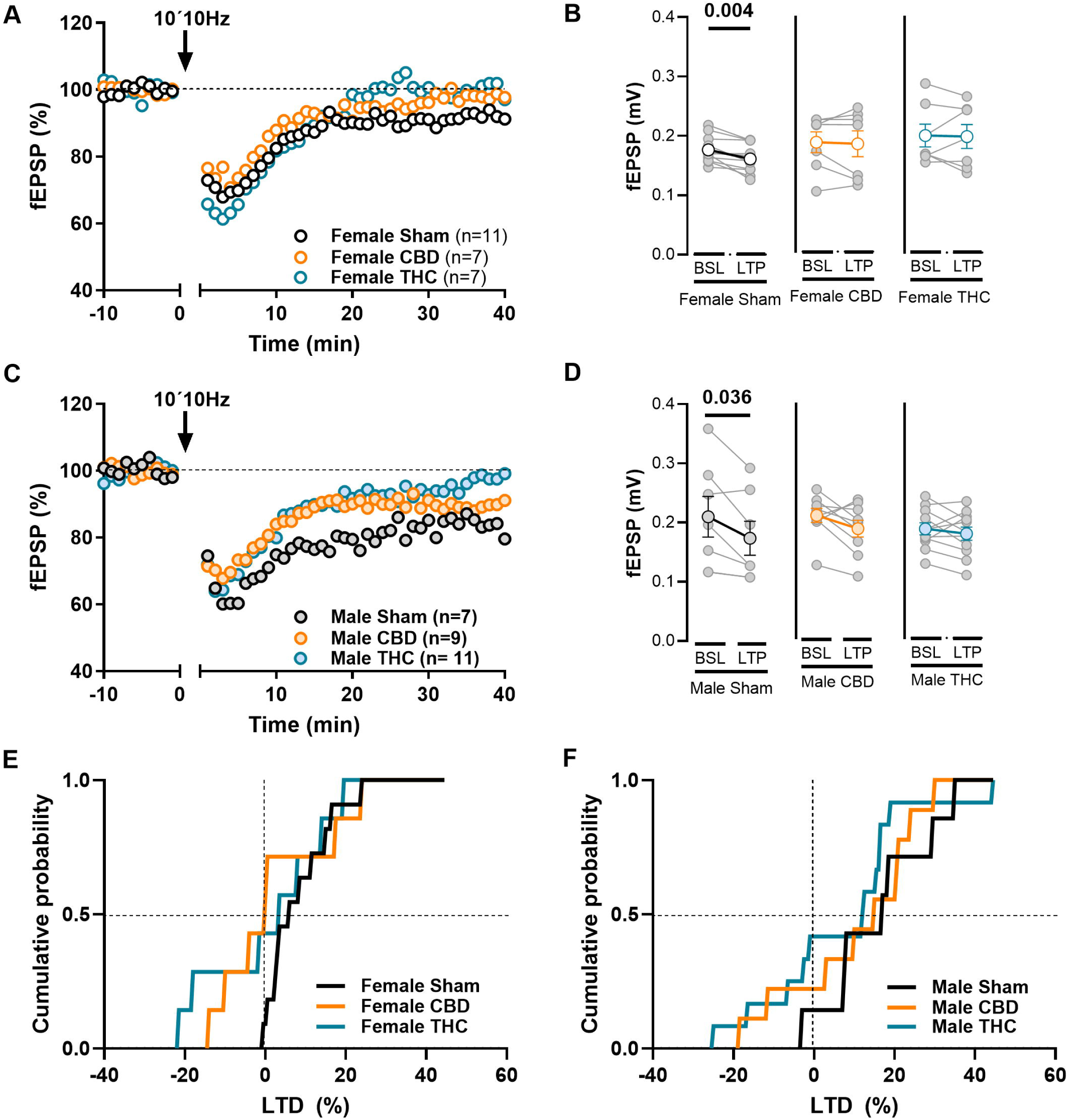
Prenatal cannabinoid exposure abolishes endocannabinoid-mediated long-term depression (eLTD). (A–C) Representative time courses of mean fEPSP amplitude before and after tetanic stimulation (10 × 10 Hz; arrow). (B–D) Individual experiments (grey dots) and group averages (colored circles) showing absolute fEPSP amplitudes during baseline (–BSL–, – 10 to 0 min) and after stimulation (–LTD–, 30–40 min post-stimulation). Tetanic stimulation induces robust eLTD in Sham females (B) and Sham males (D), whereas this form of synaptic plasticity is completely abolished in adult offspring prenatally exposed to CBD or THC. (E, F) Under control conditions, LTD is reliably induced in both sexes. In contrast, prenatal cannabinoid exposure abolishes LTD, with a substantial proportion of offspring instead exhibiting a potentiation response. Data are presented as mean ± SEM and were analyzed using the paired Wilcoxon test. Only statistically significant differences are indicated (*p < 0.05). (E, F) Cumulative probability plots show the percentage of induced LTD; negative values indicate LTP outcomes.

Importantly, the loss of eLTD in THC-exposed female mice contrasts with our earlier findings in rats, where this form of plasticity was selectively preserved in females^10^. In mice of both sexes (**Fig. 7E, F**), however, prenatal exposure to either cannabinoid produced an equivalent outcome: CBD was just as effective as THC in abolishing eLTD, indicating that CBD does not confer protection against this form of synaptic disruption. Thus, both cannabinoids converge on a shared functional endpoint in the mouse mPFC: the complete loss of activity-dependent synaptic depression—even in females

### Prenatal CBD, but not THC, inhibits prefrontal LTP

To assess whether the deficits in synaptic flexibility extended to excitatory potentiation, we applied theta-burst stimulation (TBS) to induce long-term potentiation (LTP), a major form of synaptic strengthening frequently altered in the PFC of models of neuropsychiatric disorders^18,23,26,27,27,28^. TBS reliably induced LTP in both Sham, CBD and THC-exposed offspring of both sexes, with fEPSP amplitudes significantly elevated above baseline (**Fig. 8**). Overall, neither prenatal CBD nor THC exposure affected LTP in females (**Fig. 8A, B, E**). In contrast, males exposed to CBD—but not THC—exhibited a marked reduction in LTP amplitude relative to their Sham counterparts (**Fig. 8C, D, F**). These findings indicate that prenatal CBD exposure restricts the dynamic range of excitatory strengthening in the male mPFC, whereas the mechanisms supporting LTP remain largely resilient to gestational THC interference in both sexes.

**Figure 8:**
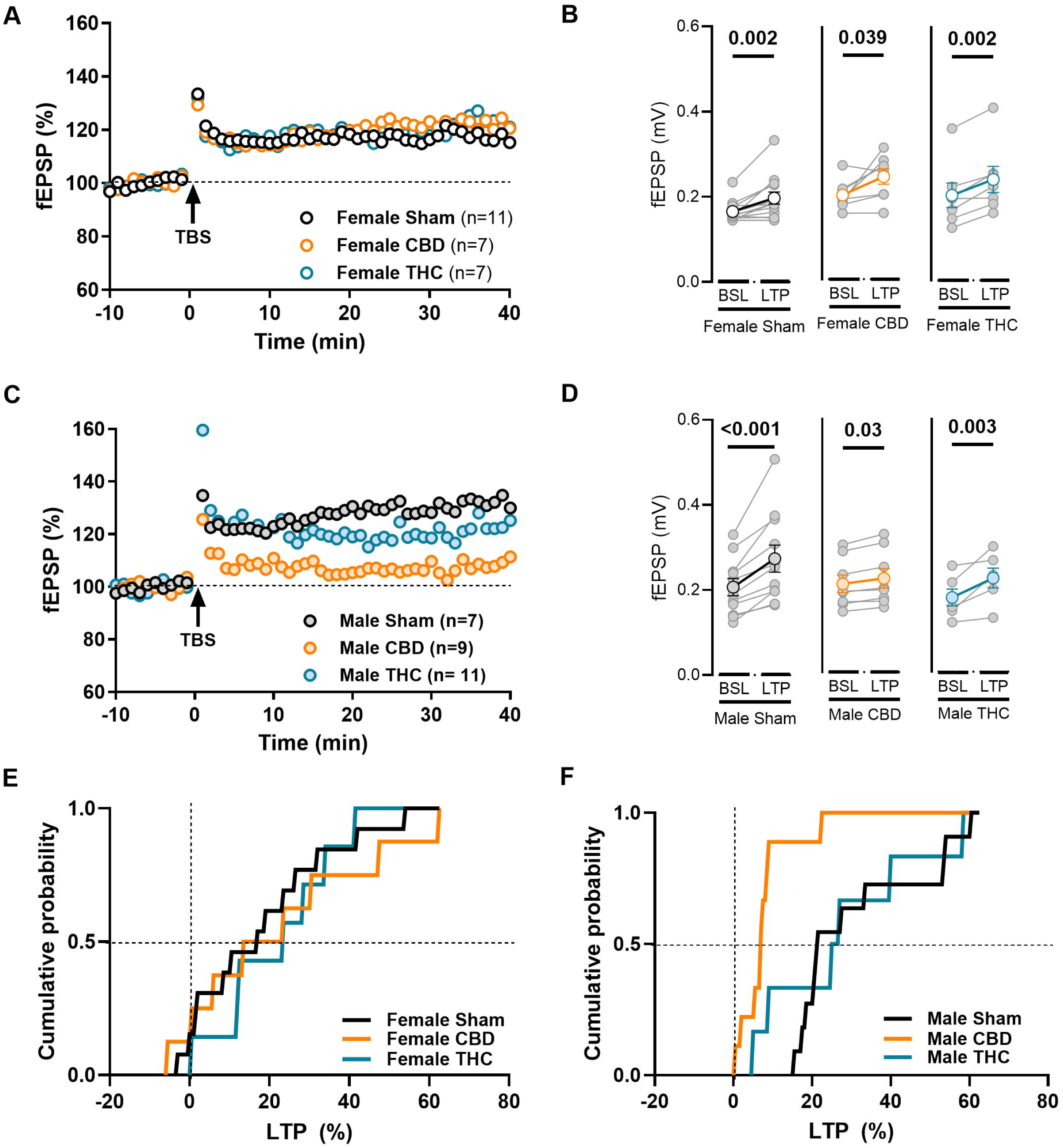
Prenatal cannabinoid exposure impairs long-term potentiation specifically in CBD-exposed males. (A, C) Representative time courses of normalized mean fEPSP amplitude before and after theta-burst stimulation (TBS). (B, D) Paired comparisons of mean absolute fEPSP amplitude during the 10-min baseline (BSL) and the 30–40 min post-stimulation period (LTP). Individual experiments (grey dots) and group averages (colored circles) are shown. TBS reliably induces LTP across treatments in females (B) and in males exposed to Sham or THC (D). (E) Female offspring exposed to cannabinoids exhibit robust LTP with amplitude increases comparable to controls. (F) In contrast, CBD-exposed males show a marked reduction in LTP magnitude, with potentiation never exceeding ∼20%. Data are presented as mean ± SEM and were analyzed using the paired Wilcoxon test. Only statistically significant differences are indicated (*p < 0.05). (E, F) Cumulative probability plots illustrate the percentage of induced LTP.

This pattern reveals a clear divergence in the “plasticity signatures” of prenatal CBD and THC. Although both cannabinoids abolish eLTD, only CBD exposure diminishes the magnitude of LTP, and this effect is specific to male offspring. Together with the eLTD data, these results suggest that in males, prenatal CBD induces a more pervasive form of synaptic rigidity by impairing the circuit’s capacity to both strengthen and weaken its connections, whereas THC selectively targets LTD.

### Sex-specific effects of gestational CBD on AMPA/NMDA ratio and NMDAR kinetics

The selective loss of LTP in male CBD-exposed progeny prompted a search for a possible cellular substrate. First, basal synaptic properties were compared by constructing input-output curves of fEPSP amplitudes against increasing stimulation intensities in the mPFC. No significant differences in the stimulus-response relationship or maximal fEPSP amplitude were found in either females (**Fig. 9A**) or males (**Fig. 9B**). This suggests that CBD treatment does not broadly alter gross synaptic recruitment or axonal excitability.

**Figure 9:**
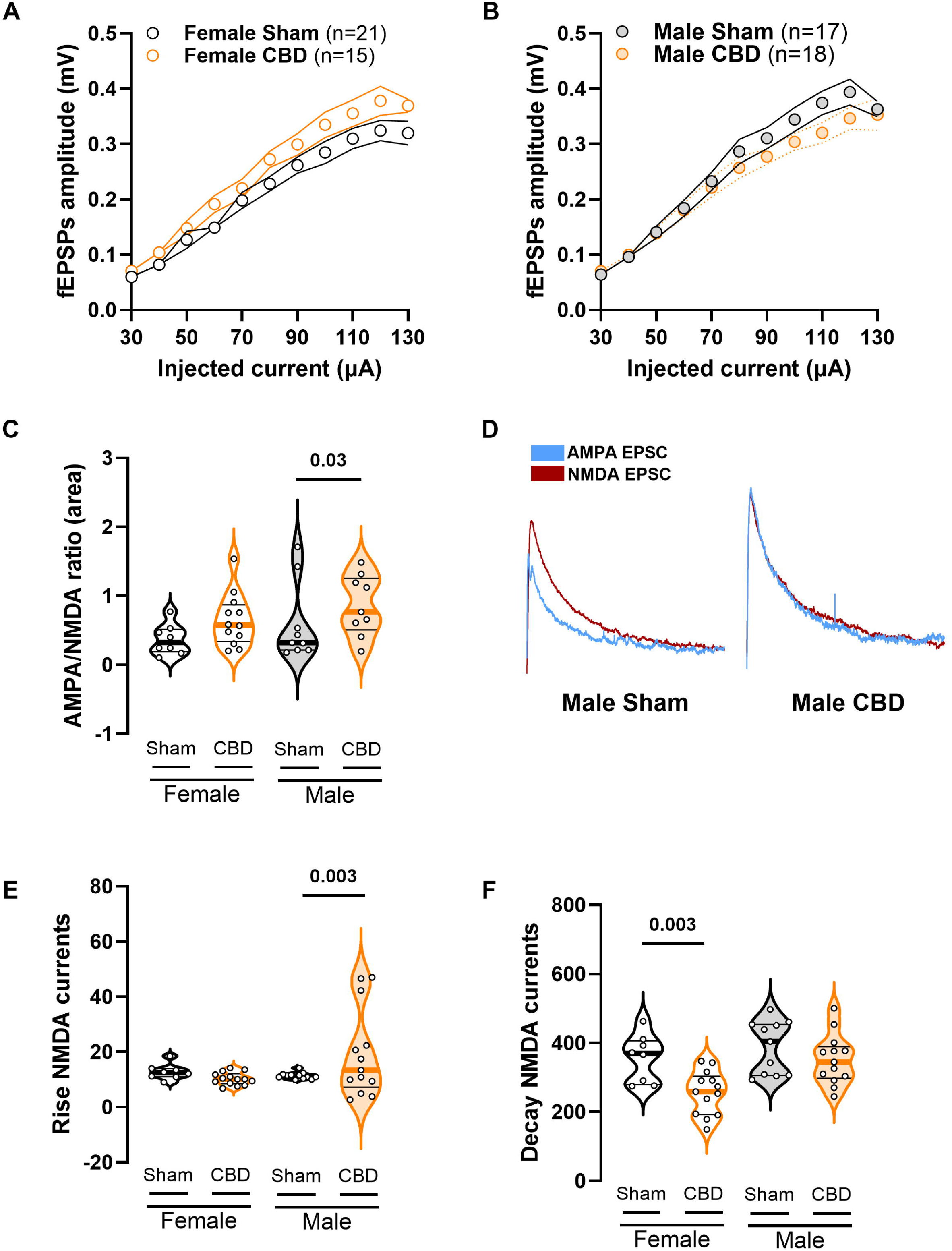
Selective increase in the AMPA/NMDA ratio in CBD-exposed male offspring. (A, B) Averaged fEPSP amplitudes plotted as a function of stimulus intensity. Synaptic strength remains stable across treatments, indicating that baseline recruitment does not account for the observed plasticity differences. (C) CBD-exposed males show a significant increase in the AMPA/NMDA ratio in layer V pyramidal neurons of the mPFC. (D) Representative traces of NMDA-EPSCs (red) evoked at +30 mV in the presence of 50 µM NBQX, and AMPA-EPSCs (blue) obtained by digital subtraction of the NMDA component from the dual-component current at +30 mV. (E, F) CBD alters the kinetics of evoked NMDA currents. Male CBD-exposed offspring exhibit slower rise times, whereas CBD-exposed females display faster decay kinetics. (A, B) Each point represents the group mean at the corresponding stimulus intensity; data are shown as mean ± SEM in XY plots and analyzed using a multiple-measures Mann–Whitney test. (C, E, F) Each point represents a single neuron; data are presented as violin plots (median and 25th–75th percentiles) and analyzed using two-way ANOVA followed by Šídák’s multiple-comparison test. Only statistically significant differences (p < 0.05) are indicated.

At excitatory synapses, the relative levels of AMPAR and NMDAR (the A/N ratio) provide a measure of synaptic integration and plasticity. This ratio serves as a proxy for synaptic strength, where an increase typically indicates the "unsilencing" of synapses or the recruitment of additional AMPA receptors into the postsynaptic membrane, a hallmark mechanism of LTP^28,29^. We hypothesized that the lack of LTP in CBD-exposed males might result from a baseline increase in the A/N ratio, creating a "ceiling effect" that impedes further potentiation. While no significant changes were observed in female mice, CBD-treated males exhibited a significant increase in the A/N ratio compared to their sham counterparts (**Fig. 9C-D**).

To further understand the underlying changes in NMDAR function, we analyzed the kinetic properties of isolated NMDA currents. In male mice, gestational CBD treatment was associated with a significant increase in the rise time of NMDA currents (**Fig. 9E**), indicating slower receptor activation or changes in subunit composition. Conversely, CBD-exposed females displayed a significant decrease in NMDA decay time (**Fig. 9F**), with no such change observed in males. Because LTP requires a fast, high-amplitude NMDAR-mediated Ca2^+^influx, the slower rise time in males may directly weakens the receptor’s ability to generate the sharp activation needed for potentiation. Thus, it is possible that this kinetic slowdown, combined with an already elevated A/N ratio, places male synapses in a state that is both less responsive to glutamate and functionally saturated at baseline, making LTP induction impossible.

## Discussion

Prenatal exposure to cannabinoids produced a profound and enduring reorganization of mPFC function, revealing a landscape of vulnerability that is both sex-specific and compound-specific. Although CBD and THC ultimately converged on a shared functional deficit, the complete loss of eLTD, the developmental trajectories leading to this endpoint were strikingly divergent. These findings demonstrate that the mPFC does not respond to prenatal cannabinoid exposure through a uniform mechanism; instead, it undergoes a selective rewiring of intrinsic excitability, synaptic strength, temporal kinetics, and plasticity that depends critically on the sex of the offspring and the identity of the cannabinoid.

The data show that the intrinsic physiology of mPFC pyramidal neurons was selectively reshaped rather than globally disrupted. THC exposure increased rheobase in females while sparing resting membrane potential, suggesting a shift toward reduced intrinsic excitability that requires stronger synaptic drive to trigger firing. In males, THC produced the opposite phenotype: intrinsic hyperexcitability driven by enhanced repetitive firing capacity, emerging independently of threshold mechanisms. CBD induced yet another pattern, selectively altering intrinsic properties in females while leaving males largely unaffected. These compound- and sex-specific signatures indicate that prenatal cannabinoids do not simply bias the circuit toward excitation or inhibition; instead, they recalibrate the rules governing neuronal responsiveness in ways that reflect cannabinoid-specific perturbations of distinct developmental programs in the male and female mPFC.

Synaptic transmission and the resulting E/I balance were reorganized along divergent lines. In females, CBD exposure produced a coordinated amplification of both excitatory and inhibitory drive, reflected in increased amplitudes, frequencies, and a rightward shift in event distributions—suggesting a homeostatic "scaled-up" architecture that preserves E/I balance despite increased throughput. In contrast, THC exposure in females led to enhanced excitatory gain paired with a large reduction in inhibitory charge transfer, resulting in a collapse of the net E/I balance toward a pro-excitatory state. In male offspring, THC reduced inhibitory transmission but did not cause a disinhibition phenotype, while CBD produced minor changes without significantly affecting synaptic strength. These divergent consequences were captured by our E/I index, which confirmed that THC significantly elevated the ratio in females, while CBD preserved balance across sexes. These findings underscore that E/I balance is an emergent property of the circuit; the female-specific collapse after THC exposure may represent a latent vulnerability that, while masked at baseline, becomes functionally consequential during plasticity-inducing challenges.

Temporal kinetics provided an additional layer of mechanistic insight into this reorganization. THC accelerated excitatory rise times across sexes, consistent with changes in AMPA receptor subunit composition or synaptic microarchitecture. Inhibitory kinetics were even more heterogeneous: CBD accelerated both rise and decay in females, whereas THC prolonged inhibitory decay in females but slowed both phases in males. These kinetic signatures indicate that prenatal cannabinoids reshape not only synaptic strength but also the temporal filtering properties of mPFC microcircuits. Such shifts in kinetics have significant potential consequences for spike timing, integration windows, and oscillatory coupling, further differentiating how each compound recalibrates the temporal dynamics of prefrontal processing.

Despite their divergent effects on intrinsic and synaptic physiology, both CBD and THC abolished eLTD in the adult mPFC. This convergence is particularly notable given that eLTD was previously preserved in THC-exposed female rats, suggesting species-specific differences in developmental sensitivity. The complete loss of eLTD across sexes and compounds indicates that endocannabinoid-dependent plasticity is a core developmental target of prenatal cannabinoid exposure, potentially reflecting disrupted maturation of CB1R signaling, interneuron recruitment, or presynaptic release machinery. Converging preclinical work shows that developmental THC exposure disrupts CB1R-dependent regulation cortical wiring ^30–34^. Complementary translational studies highlight that such molecular and circuit-level perturbations map onto structural and functional changes in human prefrontal cortex, particularly when cannabis exposure occurs during sensitive developmental windows ^3,35^.

CBD produced an additional deficit not seen with THC: a selective impairment of LTP in males. Mechanistically, this lack of LTP was accompanied by a significant increase in the AMPA/NMDA ratio, compatible with the idea that synapses have undergone a baseline potentiation prior to any plasticity-inducing stimulus, thereby occluding further LTP ^36^. In CBD-exposed males, this occlusion is further compounded by the observed increase in NMDAR rise time, which likely reduces the rate of influx below the threshold required to activate the signaling cascades necessary for potentiation. Interestingly, CBD-exposed females exhibited a distinct kinetic phenotype characterized by accelerated NMDAR decay times without the concomitant increase in basal synaptic strength seen in males. While a faster decay typically limits the total charge transfer, the lack of a "ceiling effect" in females may explain why their capacity for LTP remains preserved.

Together with the universal loss of eLTD, this reduction in potentiation creates a bidirectional synaptic rigidity in which circuits can neither strengthen nor weaken their connections; such rigidity is likely to impair cognitive flexibility, working memory and adaptive decision-making, functions that depend critically on mPFC plasticity ^37,38^. This interpretation is also consistent with our recent behavioral observations in the same treatment groups, where both cannabinoids increased repetitive defensive burying and CBD selectively heightened risk-assessment postures in females^13^. By contrast, THC selectively targeted LTD while sparing LTP, indicating that the two cannabinoids disrupt distinct components of the plasticity machinery.

Together, these findings support a mechanistic model in which prenatal cannabinoids induce sex-specific and compound-specific rewiring of the mPFC. THC primarily disrupts inhibitory control, producing hyperexcitability in males and a collapse of E/I balance in females. CBD induces a coordinated scaling of synaptic inputs in females and a rigidification of plasticity mechanisms in males. Despite these divergent pathways, both cannabinoids converge on the loss of eLTD, revealing a shared vulnerability of endocannabinoid-dependent maturation. This framework highlights that prenatal cannabinoid exposure does not simply dampen or excite the developing cortex; it reprograms the rules of synaptic integration and plasticity in a manner that depends on both sex and compound identity.

These results carry important translational implications. CBD is widely perceived as benign ^9^, yet our data reveal that it can induce synaptic rigidity and impair LTP in males, while THC produces destabilizing shifts in E/I balance in females. The sex-specificity of these effects underscores the need for sex-disaggregated analyses in both preclinical and clinical studies. Moreover, the convergence on eLTD loss suggests that even moderate prenatal cannabinoid exposure may compromise the maturation of prefrontal circuits essential for executive function, emotional regulation, and cognitive flexibility. By revealing the mechanistic diversity and convergence of cannabinoid-induced developmental perturbations, this work provides a framework for understanding the long-term cognitive risks associated with prenatal cannabinoid exposure and highlights the need for caution regarding cannabinoid use during pregnancy.

## Author Contributions

A.C.-R.: conceptualization, data curation, formal analysis, validation, writing—review and editing. D.I.: data curation, writing—review and editing. O.L.: D.I.: data curation. S.W: formal analysis; P.C.: conceptualization, methodology, project administration, supervision. O.J.J.M.: conceptualization, supervision, funding acquisition, methodology, project administration, writing—original draft, review, and editing. All authors have read and agreed to the published version of the manuscript.

## Acknowledgements

The authors are grateful to the Chavis-Manzoni team members for helpful discussions. The authors would like to acknowledge the use of the large language model Gemini (Google) for structural and linguistic refinements.

