## Supplemental data 1 for "Compound- and sex-specific medial prefrontal cortex rewiring after prenatal THC and CBD exposure"

**Supplementary File 1: Codes in R**

***Kolmogorov-Smirnov (KS) statistics***

*# Load necessary library*

*if (!require("dplyr")) install.packages("dplyr")*

*library(dplyr)*

*# 1. Load the data*

*# Replace "data.csv" with the path to your spreadsheet df <- read.csv("data.csv")*

*# 2. Define a helper function to run the KS test*

*run_ks_test <- function(test_col, sham_col, sex, treatment, sector) {*

*# Extract values and remove NAs*

*test_vals <- na.omit(df[[test_col]])*

*sham_vals <- na.omit(df[[sham_col]])*

*# Perform the two-sample Kolmogorov-Smirnov test*

*ks_res <- ks.test(test_vals, sham_vals)*

*# Return results as a data frame row*

*return(data.frame(*

*Sex = sex,*

*Comparison = treatment,*

*Sector = sector,*

*D_statistic = round(as.numeric(ks_res$statistic), 3),*

*p_value = round(as.numeric(ks_res$p.value), 4)*

*))*

*}*

*# 3. Execute all comparisons*

*ks_table <- rbind(*

*# Female Comparisons*

*run_ks_test("Female_CBD_E", "Female_Sham_E", "Female", "CBD vs Sham", "Excitatory"),*

*run_ks_test("Female_CBD_I", "Female_Sham_I", "Female", "CBD vs Sham", "Inhibitory"),*

*run_ks_test("Female_THC_E", "Female_Sham_E", "Female", "THC vs Sham", "Excitatory"),*

*run_ks_test("Female_THC_I", "Female_Sham_I", "Female", "THC vs Sham", "Inhibitory"),*

*# Male Comparisons*

*run_ks_test("Male_CBD_E", "Male_Sham_E", "Male", "CBD vs Sham", "Excitatory"),*

*run_ks_test("Male_CBD_I", "Male_Sham_I", "Male", "CBD vs Sham", "Inhibitory"),*

*run_ks_test("Male_THC_E", "Male_Sham_E", "Male", "THC vs Sham", "Excitatory"),*

*run_ks_test("Male_THC_I", "Male_Sham_I", "Male", "THC vs Sham", "Inhibitory")*

*)*

*# 4. View the final table*

*print(ks_table)*

*# Optional: Save to CSV*

*# write.csv(ks_table, "KS_Results_Table.csv", row.names = FALSE)*

***Non-Parametric Bootstrap E/I Index***

*set.seed(28)*

*# 1 Load data*

*df <- read.csv('data.csv') # Read cellular E and I values*

*# 2 Define contrasts of interest*

*bg_contrasts <- list(c('Female Sham', 'Male Sham'),*

*c('Female Sham', 'Female CBD'),*

*c('Female Sham', 'Female THC'),*

*c('Female CBD', 'Female THC'),*

*c('Male Sham', 'Male CBD'),*

*c('Male Sham', 'Male THC'),*

*c('Male CBD', 'Male THC'))*

*# 3 Bootstrap "neural populations" to calculate E/I index*

*groups <- unique(df$Group)*

*B <- 10000*

*df_B <- data.frame(Group=rep(groups, each=B), EII=rep(0, length(groups)*B))*

*# Bootstrap and calculate E/I index*

*for (i in 1:length(groups)){ # per group*

*this_group <- groups[i]*

*for (j in 1:B){	# per iteration*

*# Average E values*

*df_E <- df[df$Group==this_group & df$Type=='E',]*

*nE <- length(df_E$Value) #round(length(df_E$Value) * 0.8)*

*Es <- sample(df_E$Value, nE, replace=TRUE)*

*E_avg <- mean(Es)*

*# Average I values*

*df_I <- df[df$Group==this_group & df$Type=='I',]*

*nI <- length(df_I$Value) #round(length(df_I$Value) * 0.8)*

*Is <- sample(df_I$Value, nI, replace=TRUE)*

*I_avg <- mean(Is)*

*# E/I Index*

*EII <- E_avg / (E_avg + I_avg)*

*df_B$EII[(i-1)*B+j] <- EII*

*}*

*}*

*# 4 Calculate between-group difference in E/I index*

*for (i_contrast in bg_contrasts){*

*df_ <- df_B[df_B$Group %in% i_contrast,]*

*EII <- df_$EII[df_$Group==i_contrast[2]] - df_$EII[df_$Group==i_contrast[1]]*

*b0 <- mean(EII < 0) * 100 # b0 % of EII below 0*

*print(sprintf('For the contrast %s - %s, %.2f%% below 0', i_contrast[2], i_contrast[1], b0))*
